# Tissue-Specific and NF-kappaB-Independent Xist RNA Localization Patterns in Female Intestinal, Blood, and Muscle Progenitors

**DOI:** 10.1101/2025.08.30.673230

**Authors:** Claudia D. Lovell, Isabel Sierra, Emma M. Welter, Donavon Sandoval-Heglund, Ning Li, Ashley Vanderbeck, Christopher J. Lengner, Christoph Lepper, Montserrat C. Anguera

## Abstract

X-Chromosome Inactivation (XCI), driven by the expression of Xist RNA and the enrichment of various repressive epigenetic marks, results in the formation of a mostly inactive X chromosome (Xi) to equalize X-linked gene expression between sexes. Unexpectedly, in unstimulated female lymphocytes that are largely quiescent, these epigenetic features are largely absent but are restored by NF-κB signaling following their activation. To determine whether these epigenetic phenotypes correlate with quiescence or NF-κB activation in other tissues, we evaluated female progenitor and stem cells from intestine, blood, and muscle. Despite known NF-κB activation, intestinal progenitors have variable Xist RNA patterns, whereas blood progenitors and neutrophils show a strong correlation between NF-κB activation status and Xist RNA localization. In contrast, muscle satellite cells (SCs) and myoblasts exhibit Xist RNA accumulation at the Xi without NF-κB activation. Xist RNA localization patterns in SCs change with age, yet adult SCs have an Xi that is mostly transcriptionally silent while allowing expression of muscle-specific X-linked genes including *Dmd*. These findings reveal that female somatic cells employ diverse, tissue-specific epigenetic mechanisms to maintain X chromosome inactivation, enabling cell type specific Xi gene expression while preserving chromosome-wide silencing.

## INTRODUCTION

Female mammals utilize X-Chromosome Inactivation (XCI) for equilibrating X-linked gene expression between the sexes. XCI initiation during early development results in the formation of a mostly inactive X chromosome (Xi) that expresses the long-noncoding RNA Xist and is enriched with various repressive histone tail modifications including H3K27me3 and H2AK119-ubiquitin (Ub), histone variant macroH2A, and DNA methylation^1–4^. The enrichment of some of these epigenetic modifications on the Xi are cytologically visible in female fibroblasts and *in vitro* differentiated pluripotent stem cells, with Xist RNA transcripts exhibiting a bright ‘cloud’ structure that overlaps the Xi territory and co-localizes with immunofluorescent enrichment of H3K27me3 and H2AK119-Ub. While it was generally assumed that all mature female somatic cells would display similar epigenetic features of XCI, numerous studies now suggest that this is not the case. For example, somatic female cells including lung progenitor alveolar type 2 (AT2) cells^5^ as well as natural killer cells and dendritic cells of the immune system^6^ have pinpoints of Xist RNA FISH that vary in size, while bone marrow derived macrophages and plasmacytoid dendritic cells lack Xist RNA ‘clouds’ or pinpoints^6^. Pre-XCI pluripotent stem cells^7^, human primordial germ cells^8^, and human and monkey preimplantation embyros^9^ exhibit dispersed patterns of XIST/Xist RNA. Unstimulated mouse and human T and B cells lack cytological enrichment of XIST/Xist RNA and heterochromatic modifications on the Xi^10–14^, yet despite this lack, the Xi is mostly transcriptionally silent with H3K27me3 and DNA methylation imprints chromosome-wide^13,15^. Surprisingly, in *vitro* stimulation of T and B cells using various antigenic stimuli results in cytological enrichment of Xist RNA and H3K27me3 and H2AK119-Ub marks at the Xi^10–14^. Notably, genetic or chemical inhibition of NF-kB nuclear translocation prevents Xist RNA tethering at the Xi^13^, providing evidence that epigenetic mechanisms regulating XCI may depend upon NF-kB signaling.

XCI maintenance, through enrichment of epigenetic modifications across the Xi, persists with each cell division into adulthood. While the majority of X-linked genes are transcriptionally silent on the Xi, there are X-linked genes that ‘escape’ XCI and are expressed from both the Xi and the active X chromosome (Xa)^16–22^. Some XCI escape genes with a Y-linked homolog, referred to as X-Y pairs, have cellular housekeeping functions including regulation of chromatin (i.e. *Kdm6a/Kdm6b; Kdm5c/Kdm5d*) and transcription (*Zfx, Zfy*). Importantly, XCI escape genes vary by both cell type and cellular activation state, as naïve and stimulated B and T cells share some XCI escape genes yet also express unique Xi-linked genes^13,23^. Surprisingly, despite the lack of Xist RNA, H3K27me3, or H2AK119Ub cytological enrichment in female B and T cells and AT2 cells, the Xi is mostly transcriptionally silent, with only 10% to 20% of X-linked genes expressed from the Xi ^5,13,23^. This frequency compares favorably with female mouse embryonic fibroblasts, which bear a canonical Xist RNA ‘cloud’, and in which 9% of expressed X-linked genes have escaped silencing^23^. Thus, while early studies suggested that cytological enrichment of Xist RNA and heterochromatic histone marks in female somatic cells would correlate with the extent of XCI escape genes, alternative mechanisms that preserve transcriptional silencing on the Xi must also exist to maintain XCI.

Quiescent naïve B and T lymphocytes residing in G0 of the cell cycle^24,25^ have a more compact nucleus, globally reduced transcription, and lack cytological enrichment of XIST/Xist RNA and heterochromatic modifications on the Xi compared to activated lymphocytes^26^. Quiescence is a unique cellular state utilized predominately by adult stem cells as a protective maintenance mechanism to allow for effective tissue homeostasis^27,28^. Some adult stem cells under steady-state conditions can exist in a quiescent state, including facultative stem cells of the intestinal epithelium^29^, adult hematopoietic stem cells (HSCs) in the bone marrow^30,31^, and satellite cells (SCs) in skeletal muscle^32^. At the cellular level, quiescence is often characterized by cells that have exited the cell cycle, exhibit low metabolic output, transcription and translation rates^33^. This contrasts with other cellular states that feature irreversible cell cycle exit such as terminal differentiation or senescence. There is evidence that female adult stem cells function with different efficiencies compared to male adult stem cells. Sex differences with proliferation and regenerative efficiency have been observed in ISCs of the small intestine^34^, HSCs^35^, and SCs of muscle^36–38^, yet the biological contributions of sex hormones and sex chromosomes for these sex differences is not well understood. Estrogen is responsible for some of these observed differences^35,39,40^, yet evidence suggests that the X-chromosome also plays a role^34^. For example, there are sex differences with skeletal muscle fiber types and sizes, with ∼4000-6000 differentially expressed mRNAs in muscle types that cluster by sex, and the genes with greatest sex differences are X-linked and lack estrogen responsive elements^41^. The XCI escape gene *Kdm6a* is one of the most highly expressed genes in female muscle, which can influence gene expression across the genome through its H3K27me3 demethylase activity and also through non-catalytic association with KMT2D to mediate p300 recruitment to enhancers^42^. Thus understanding how XCI is maintained in adult stem cells and the X-linked genes expressed from the Xi can reveal genetic contributions underlying sex differences with proliferation and regenerative pathways mediated by stem cells.

A defining feature of quiescent cells is their ability to re-enter the cell cycle and re-initiate proliferation and cell division in response to injury or various signals^27,43^. NF-κB activation is required to regulate HSC homeostasis and function in multiple cell types^44^. Adult stem cells respond to external stimuli or injury by re-entering the cell cycle and differentiating into specific cell types and can also self-renew. NF-κB signaling is also observed in Lgr5+ crypt base columnar stem cells (which are actively cycling), Paneth cells, and also the ‘+4/+5’ position secretory progenitor cells, and NF-κB is required for Paneth vs goblet cell differentiation, cellular homeostasis, and post-injury regeneration^45,46^. In skeletal muscle, canonical NF-κB signaling promotes myoblast proliferation and negatively regulates their differentiation, while chronic NF-κB activation in SCs results in telomere shortening and severe skeletal muscle defects^47,48^. Thus, activated NF-κB signaling is necessary for some, but not all, types of adult stem cells.

Here we asked how NF-κB activation correlates with Xist RNA localization at the Xi across female adult stem cells from various tissues. We found that NF-κB activation status is usually positively correlated with the presence of Xist RNA clouds for most adult stem and progenitor cells in the intestine, blood, and terminally differentiated immune cells, yet some progenitor cells exhibited dispersed Xist RNA localization patterns or completely lacked detectable Xist RNA signals. Surprisingly, SCs and myoblasts from muscle exhibited a negative correlation with NF-κB activation and Xist RNA clouds, and express genes important for muscle function from the Xi. Our findings indicate that female somatic cells use diverse, tissue-specific epigenetic strategies to maintain XCI, allowing selective Xi gene expression while preserving overall chromosome silencing.

## RESULTS

### Xist RNA localization patterns are variable across stem cells and progenitor cells of the intestinal crypt compartment

While most female somatic cells exhibit a cytological Xist RNA ‘cloud’^49^, quiescent naïve T and B cells, that have NF-κB retained in the cytoplasm, lack Xist RNA clouds. Upon immune cell stimulation and NF-κB activation, Xist RNA clouds coat the Xi^10–12^. We asked whether Xist RNA clouds correlate with NF-κB activation in adult stem and progenitor cells, and particularly other quiescent cell types outside of the immune system. First, we investigated Xist RNA localization in small intestinal crypts, which contain intestinal stem cells (ISCs) and progenitors ^50^, and where most cells have active NF-κB signaling^45^. We performed Xist RNA FISH using fixed tissue sections of intestinal crypts derived from the jejunum^51^. We quantified the frequency of cells that contained a localized Xist RNA cloud at each position of the crypt from multiple female mice. We observed that fewer cells at the +4 position of the crypt, which are typically quiescent and considered to be facultative stem cells,^52–54^ have Xist RNA clouds (18-33%; ****p < 0.001) compared to the majority of cells at all other positions along the crypt with greater frequencies of canonical Xist RNA clouds (56-90%) (**Figure 1A, B**). Because cells of the lower crypt are a mixture of ISCs, Paneth cells, and progenitor cells, we investigated Xist RNA localization patterns in sorted stem cells and progenitor cells that express cell-type specific markers using female fluorescent reporter mice. Hopx is an atypical homeobox protein expressed in intestinal crypts, and a CreER reporter cassette knocked into the endogenous *Hopx* 3’UTR primarily marks slow-cycling quiescent and active cycling stem cells^55^. We treated Hopx reporter female mice (*Hopx^ERCre/+^; R26^LSL-tdTomato^*^55^) with tamoxifen, then sorted Tomato positive and negative cells from the small intestine and determined Xist RNA localization via FISH. We found that Hopx Tomato+cells, which primarily reside in G0 of the cell cycle^29^, had a higher frequency of nuclei lacking detectable Xist RNA signals and slightly lower frequencies of nuclei with localized Xist RNA patterns compared to Tomato-cells, although the differences were not statistically significant (**Figure 1C, D**). Next, we isolated intestinal cells expressing leucine-rich repeat-containing G protein 5 (Lgr5) crypt-base columnar cells, which are an actively cycling stem cell population that resides at the base of the crypt and can generate the epithelial cell lineages of the intestine^56^. Using *Lgr5^EGFP-IRES-CreERT^*^2^ female mice, we isolated both Lgr5+ and Lgr5-intestinal cells and found that the majority of Lgr5+ cells (∼60%) have robust Xist RNA clouds or dispersed RNA patterns (∼20%), almost identical to Lgr5-cells (**Figure 1E, F**). We also investigated Xist RNA localization patterns for enteroendocrine cells within the intestinal crypt which express the neuroendocrine secretory protein Chromogranin A (Chga), and these cells can become activated in response to injury^57,58^. We used tamoxifen-treated treated *Chga^CreERT2-2A-tdTomato^* female mice to isolate Chga+ and Chga-cells. We found that Chga+ cells from the intestine lacked Xist RNA clouds and were mostly devoid of bright Xist RNA patterns, unlike Chga-cells that contained bright Xist RNA clouds **(Figure 1G, H)**. Taken together, our results show that Xist RNA localization patterns are variable across stem cells and progenitor cells of the intestinal crypt compartment and that Xist RNA clouds do not correlate with the active NF-κB signaling characteristic of intestinal stem/progenitor cells^45^.

**Figure 1.**
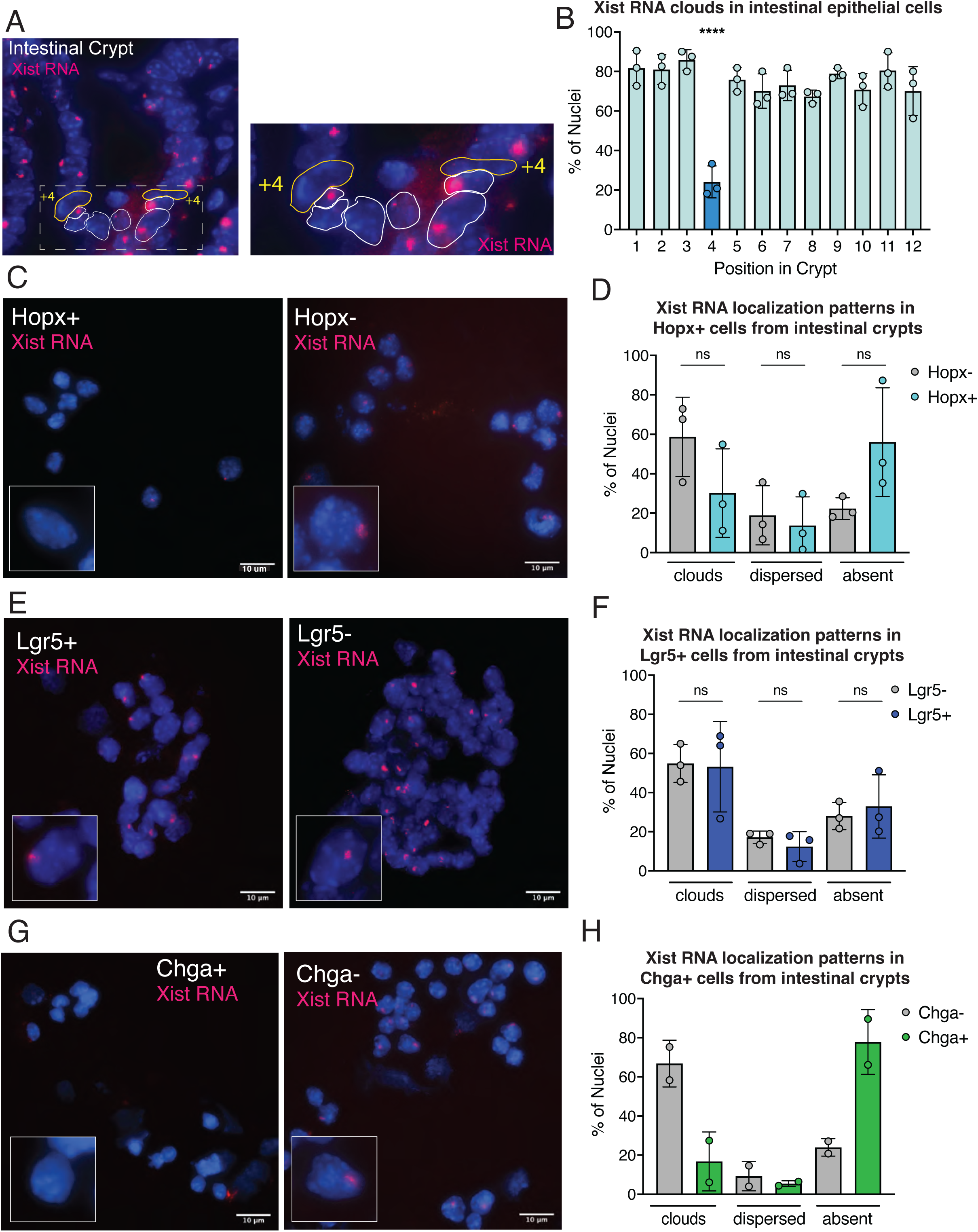
Variable Xist RNA localization patterns in stem and progenitor cells of the intestinal crypt compartment. **A.** Representative intestinal crypt from tissue section of the small intestine of female mice for Xist RNA FISH imaging. The crypt area is shown in white dashed line box, and zoomed area is shown to the right. The +4 position is shown. **B.** Quantification of percentage of nuclei with Xist RNA clouds for each of the 12 crypt positions from female mice (n=3 biological replicates of whole intestinal crypt). **** *p* < 0.0001 by ordinary one-way ANOVA followed by Dunnett’s multiple comparisons test. **C.** Representative Xist RNA FISH images of sorted Hopx+ (left) and Hopx-(right) cells of the small intestine. Inset shows zoom image of representative nucleus. **D.** Quantification of Xist RNA localization patterns (clouds, dispersed, absent) for Hopx+ (blue) and Hopx-(grey) sorted intestinal cells from female mice (n=3 mice). “Ns” not significant by unpaired t test. **E.** Representative Xist RNA FISH images of sorted Lgr5+ (left) and Lgr5-(right) cells of the small intestine. Inset shows zoom image of representative nucleus. **F.** Quantification of Xist RNA localization patterns for Lgr5+ (dark blue) and Lgr5-(grey) sorted intestinal cells from female mice (n=3 mice). “Ns” not significant by unpaired t test. **G.** Representative Xist RNA FISH images of sorted Chga+ (left) and Chga-(right) cells of the small intestine. Inset shows zoom image of representative nucleus. **H.** Quantification of Xist RNA localization patterns for Chga+ (green) and Chga-(grey) sorted intestinal cells from female mice (n=2 mice).

### Blood progenitor cells and neutrophils with NF-κB activation exhibit Xist RNA clouds, irrespective of cell cycle status

We investigated Xist RNA localization patterns in hematopoietic stem cells (HSCs) which exhibit basal NF-κB activation that is required for HSC self-renewal and quiescence^59^. HSCs (CD150+CD48-) are quiescent and slow cycling yet can undergo self-renewal to sustain the stem cell pool. HSCs can differentiate into multi-potent progenitors (MPPs) to generate differentiated immune cells, which also rely on NF-κB activation for homeostasis^59,60^. We isolated quiescent HSCs (Lin^-^Sca-1^+^cKit^+^CD150^+^CD48^-^) and actively cycling MPPs (Lin^-^Sca-1^+^cKit^+^CD150^-^Cd48^-^)^61^ from bone marrow of female mice. We found that most HSCs and MPPs have localized Xist RNA clouds yet also have a high frequency of dispersed Xist RNA patterns (**Figure 2A, B**). Thus, HSCs have canonical patterns of Xist RNA localization, which correlates with known activation of NF-κB signaling pathways in these cells, similar to *in vitro* stimulated lymphocytes that have exited quiescence and have active NF-κB signaling^13^.

**Figure 2.**
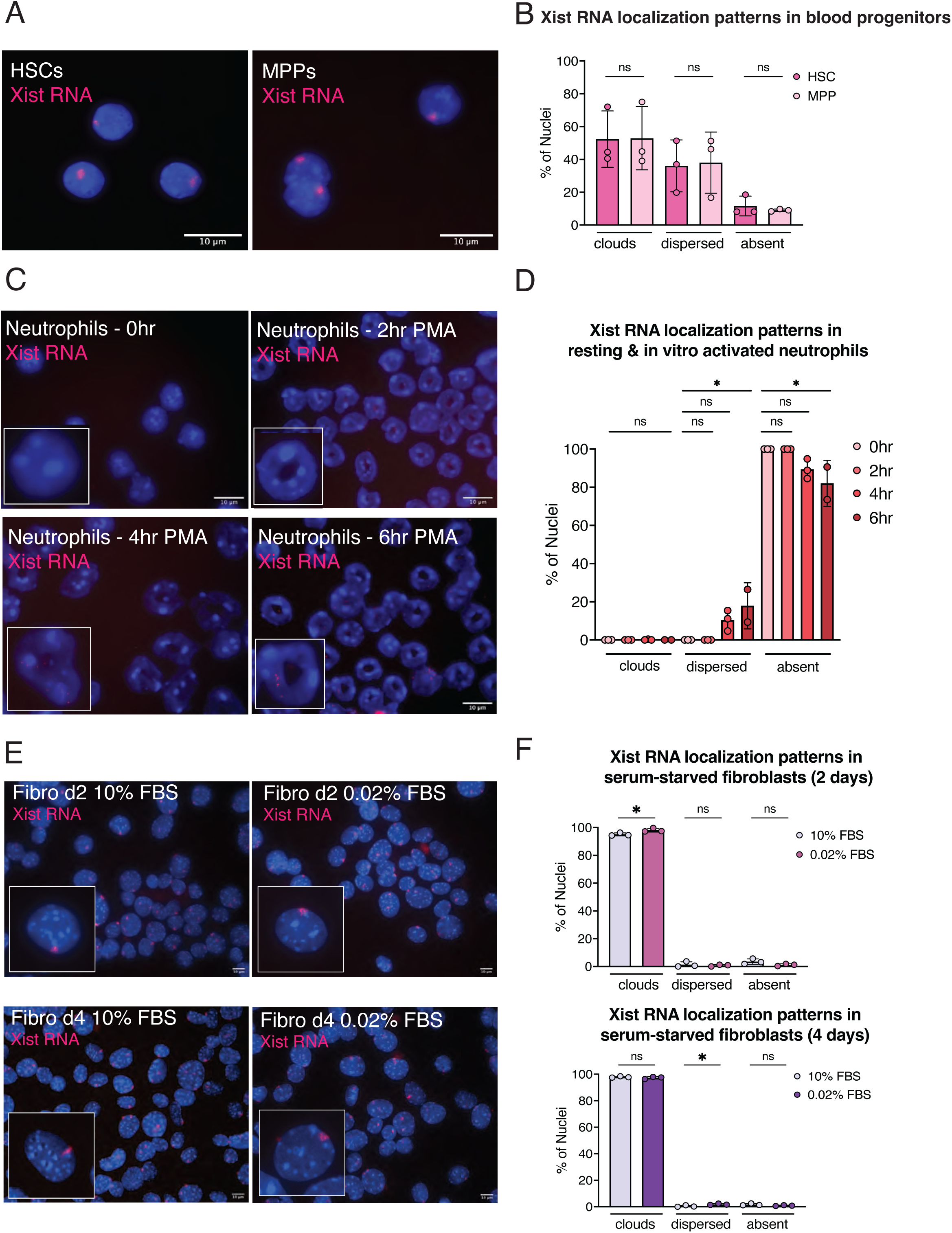
Blood progenitors, irrespective of cell cycle rate, exhibit Xist RNA clouds yet terminally differentiated neutrophils lack Xist RNA signals. **A.** Representative Xist RNA FISH images of slow cycling HSCs (Lin^-^Sca-1^+^cKit^+^CD150^+^CD48^-^) and actively cycling MPPs (Lin^-^Sca-1^+^cKit^+^CD150^-^Cd48^-^). **B.** Quantification of Xist RNA localization patterns (clouds, dispersed, absent) for HSCs and MPPs from female mice (n=3 mice). “Ns” not significant by unpaired t test. **C.** Representative Xist RNA FISH images of primary female neutrophils (“0hr”) or neutrophils cultured with PMA for 2, 4, or 6 hours. Inset shows zoom image of representative nucleus. **D.** Quantification of Xist RNA localization patterns for female unstimulated neutrophils (“0hr”), and in vitro PMA-stimulated neutrophils for 2-6 hours (n=3 biological replicates for each timepoint; n=2 for 6hr). * *p* < 0.05 by ordinary one-way ANOVA followed by Dunnett’s multiple comparisons test. **E.** Representative Xist RNA FISH images of primary female mouse embryonic fibroblasts (MEFs) cultured in low (0.02%) or replete (10%) serum (FBS) to induce cellular quiescence. Inset shows zoom image of representative nucleus. **F.** Quantification of Xist RNA localization patterns for female MEFs (n=3 biological replicates) cultured for 2 days (top) or 4 days (bottom) for each serum condition. * *p* < 0.05 by unpaired t test.

Because HSCs have Xist RNA clouds and can differentiate into naïve T and B cells which lack Xist RNA clouds, we investigated Xist RNA localization patterns in HSC lineage-derived neutrophils, which are terminally differentiated and post-mitotic innate immune cells that respond to antigenic stimulation. Resting neutrophils have reduced transcriptional activity and cytoplasmic retention of NF-κB, and neutrophil stimulation *in vitro* promotes NF-κB nuclear translocation, increased gene expression, and cytokine production^62^. We isolated resting neutrophils from bone marrow of female mice and found that naïve neutrophils completely lacked detectible Xist RNA signals in the nucleus (**Figure 2C, D**), similar to naïve mouse T and B cells^11,12,63^. Next, we stimulated neutrophils *in vitro* for 2 – 6 hours using phorbol myristate acetate (PMA), which activates the NF-κB pathway^64^, and observed the appearance of dispersed Xist RNA patterns at 4 and 6 hours post-stimulation in approximately ∼10-20% of neutrophils (**Figure 2C, D**). We did not observe localized Xist RNA clouds in 2-6hrs post-stimulation neutrophils, and the majority of cells had undergone cellular apoptosis and NETosis (neutrophil extracellular trap formation) at PMA stimulation timepoints beyond 6 hrs^65^. Thus, resting neutrophils lack localized Xist RNA clouds and PMA-induced activation results in low yet detectable dispersed Xist RNA patterns, despite the activated NF-κB state. In sum, there is variable correlation between Xist RNA localization at the Xi and NF-κB signaling status for HSCs and neutrophils of the immune system.

### Artificial induction of cell cycle exit using serum depletion does not impact Xist RNA clouds in female fibroblasts

Fibroblasts have robust Xist RNA clouds, and we wondered whether Xist RNA localization would be affected by inducing exit from the cell cycle using serum starvation^66^, which blocks fibroblast cell division and reduces NF-κB activity by reducing levels of nuclear p65, an NF-κB subunit, by 50% after 2-4 hrs^67^. We exposed primary female murine embryonic fibroblasts (MEFs) to serum starvation for 2 or 4 days and then performed Xist RNA FISH. Nearly all (∼90%) serum starved MEFs had Xist RNA cloud patterns similar to cycling fibroblasts in complete medium (**Figure 2E, F**). Yet, we noticed that serum starvation for 2 days slightly increased the percentage of nuclei with Xist RNA clouds (**Figure 2E, top**), yet serum starvation for 4 days subtly but significantly increased the frequency of nuclei with dispersed Xist RNA patterns (**Figure 2E, bottom**). We conclude that serum starvation of female MEFs, which reduces NF-κB activity and impedes cell division, does not result in high levels of dispersed Xist RNA patterns.

### Muscle satellite cells (SCs) exhibit variable patterns of Xist RNA localization that change with age, and differentiation to myoblasts increases Xist RNA and H2AK119Ub localization at the Xi

We next investigated Xist RNA localization patterns in skeletal muscle stem cells, i.e. satellite cells (SCs). During adult homeostasis, SCs are maintained in a quiescent state, yet readily respond to injury becoming activated myoblasts that enter the cell cycle to undergo several rounds of cell division, then terminally differentiate and ultimately fuse together or with injured myofibers to form newly regenerated myofibers to complete the repair process^32^. While most SC-derived myoblasts generate myonuclei for myofiber repair and regeneration, a small subset of SCs self-renew and return to quiescence, thereby maintaining the muscle SC pool. NF-κB signaling is activated in SCs and proliferating myoblasts upon injury, yet NF-κB is inhibited during differentiation and myotubule formation^68^. Xist RNA localization patterns were analyzed in freshly isolated quiescent SCs (via FACS using mice harboring a Pax7-GFP knock-in allele (*Pax7*^Avi−2A-GFP^)^69^ by FISH. We observed that, unlike stem cells of the intestinal crypt, HSCs, and neutrophils, SCs exhibit four distinct patterns of Xist RNA localization (**Figure 3A**). While most SCs had a bright pinpoint of Xist RNA signal (∼40-70%) or lacked detectable Xist RNA transcripts (∼20-40%), we did observe Xist RNA clouds in some SCs (∼15-25%) (**Figure 3A, C**). We cultured FACS-isolated SC-derived primary myoblasts, performed Xist RNA FISH, and found that nearly all myoblasts had Xist RNA clouds (∼95%) (**Figure 3B, C**).

**Figure 3.**
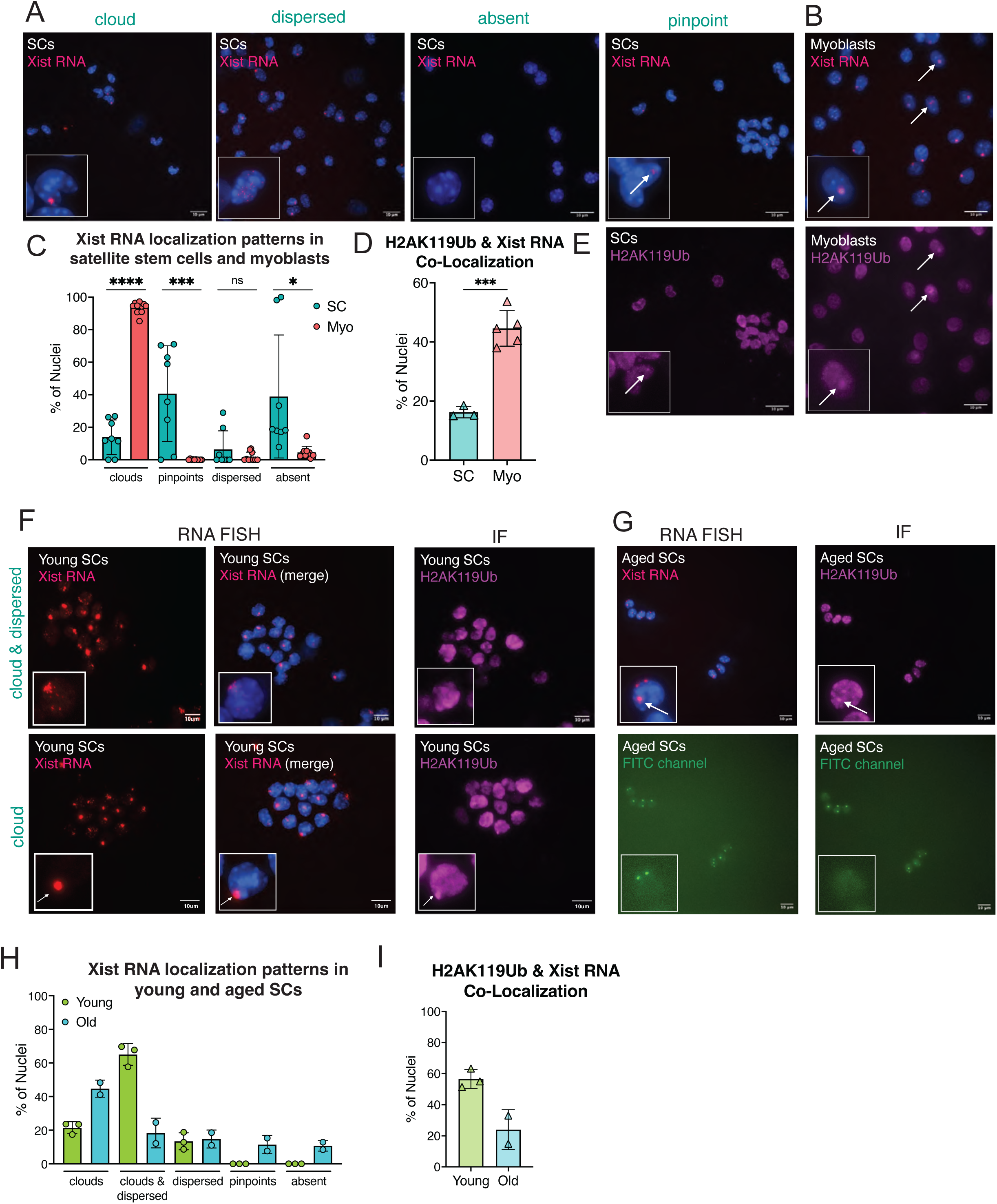
Muscle satellite cells (SCs) have variable patterns of Xist RNA localization that change across lifespan. **A.** Xist RNA FISH analyses of female SCs with representative examples of 4 localization patterns. Inset shows zoom image of representative nucleus. **B.** Sequential Xist RNA FISH followed by immunofluorescence (IF) for H2AK119-Ub for in vitro differentiated myoblasts. White arrows denote co-localization of Xist RNA signal with focus for H2AK119-Ub. Inset shows zoom image of representative nucleus. **C.** Quantification of 4 Xist RNA localization patterns in female SCs (n=8 mice) and myoblasts (n=10 mice). ** *p* < 0.01, *** *p* < 0.001, **** *p* < 0.0001 by unpaired t test. **D.** Quantification of Xist RNA and H2K119-Ub colocalization in female SCs (n=3 mice) and myoblasts (n=5 mice). *** *p* < 0.001 by unpaired t test. **E.** Representative H2AK119Ub immunofluorescence (IF) images of SCs and myoblasts. The same field is shown for ‘pinpoint’ Xist RNA FISH image in A. White arrows indicate H2AK119Ub foci that overlap with an Xist RNA signal. **F.** Sequential Xist RNA FISH and IF analyses of female SCs from postnatal day 5 (‘young’) female mice (n=3 mice) showing representative examples of Xist RNA ‘cloud’ and ‘cloud & dispersed’ patterns. White arrows indicate H2AK119Ub foci that overlap with an Xist RNA cloud by Xist RNA FISH. Inset shows zoom image of representative nucleus. **G.** Sequential Xist RNA FISH and IF analyses of female SCs from 27 month aged (‘old’) female mice (n=2 mice) showing representative examples of Xist RNA clouds. Inset shows zoom image of representative nucleus. White arrows indicate the Xi, with colocalization of both Xist RNA and H2AK119Ub focus. White arrowheads indicate non-specific (and non-overlapping with Xi) signals in both Cy3 and FITC channels during RNA FISH which were not detected during IF. **H.** Quantification of 5 Xist RNA localization patterns in female young (green; n=3) and old (blue; n=2) SCs from female mice. **I.** Quantification of Xist RNA and H2K119-Ub colocalization in female SCs from young (n=3 mice) and old (n=2 mice). Quantification of percentage of nuclei with H2AK119 foci in n=3 young SCs, n=3 adult SCs, and n=2 old SCs.

Next, we performed sequential Xist RNA FISH and IF for the heterochromatic histone tail modification H2AK119-ubiquitin (Ub) in SCs and myoblasts. H2AK119Ub forms a cytologically detectible focus localized to the Xi in most female somatic cells^70^. We quantified the percentage of nuclei exhibiting co-localization of Xist RNA and H2AK119Ub signals in SCs and *in vitro* cultured SC-derived myoblasts and observed that myoblasts have significantly higher levels of co-localization with Xist RNA and H2AK119-Ub foci (**Figure 3D, E**) compared to SCs with Xist RNA pinpoints. Some SC nuclei with Xist RNA pinpoints exhibited overlap with small foci of H2AK119Ub (**Figure 3D, E**). With aging, the regenerative capacity of SCs declines, which has been ascribed to compromised molecular regulation of SC function including the regulation of quiescence, proliferation and self-renewal^71^. We investigated Xist RNA and H2AK119Ub localization patterns in SCs from post-natal day 5 (‘young’; n=3) and 27 months (‘aged’; n=2) female mice. Surprisingly, we found that most SCs from ‘young’ mice exhibited Xist RNA clouds together with dispersed Xist RNA patterns (‘cloud & dispersed’ (**Figure 3F**)), unlike SCs from adult mice. We also observed that there were fewer SC nuclei from young mice with either Xist RNA clouds or dispersed Xist RNA patterns (**Figure 3F, H**), unlike SCs from adult female mice which exhibited pinpoints or lacked detectible Xist RNA signal. ‘Aged’ SCs, unlike adult SCs, primarily exhibited localized Xist RNA clouds (∼45%), with even distribution of the other 4 localization patterns (**Figure 3G, H**). Young SCs exhibited high levels of co-localization of Xist RNA with H2AK119Ub foci (∼60%) compared to adult (∼20%) and aged (∼20%) SCs (**Figure 3F (bottom row), I**). We observed non-specific focal staining following Xist RNA FISH and IF exclusively for ‘aged’ SCs, which did not overlap with Xi signals (**Figure 3G**, **bottom panels**). Thus, SCs have varied patterns of Xist RNA localization that change with age, and differentiation into myoblasts increases Xist RNA and H2AK119Ub localization at the Xi.

### Muscle-specific X-linked genes, including Dmd, are expressed from the Xi in SCs and myoblasts

We quantified allele-specific transcription from the Xi in SCs and myoblasts in female adult mice using an F1 mouse model of skewed XCI (**Figure 4A**). C57BL/6 female mice with a heterozygous *Xist* deletion are mated to castaneus male mice, generating female F1 mice with a castaneus-derived Xi and a C57BL/6-derived active X (Xa) chromosome lacking *Xist* expression^5,13,23^, and X-linked SNPs between strains distinguishes expression from each allele. Global gene expression in SCs and myoblasts were distinct by principal component analysis (**Figure S1A**), and allele specific reads from the maternal and paternal chromosomes were similar except for the X chromosomes in both cell types (**Figure S1B-C**). Examination of allele-specific expression patterns for X-linked *Xist* and autosomal imprinted genes *(Igfr2, Peg3)* confirmed appropriate parental expression (**Figure S1D-F**). *Pax7* was robustly expressed in SCs (**Figure S1G**) while *Myod1* was highly expressed in myoblasts (**Figure S1H**) confirming expression of cell-type specific transcripts^72,73^. We observed that the Xi is mostly transcriptionally silent compared to the Xa in both SCs and myoblasts (**Figure 4B**). Yet, 13% of expressed X-linked genes are expressed from the Xi in SCs and 11% are expressed from the Xi in myoblasts, which is similar to the frequency of XCI escape genes in B and T cells^13,23^. We found 32 X-linked genes that are expressed from the Xi in SCs and 38 X-linked genes that are expressed from the Xi in myoblasts, of which 25 are shared between cell types (**Figure 4C, D**). Remarkably, we found that *Dmd* — the gene encoding dystrophin, a structural protein essential for muscle cell function that is mutated in Duchenne Muscular Dystrophy^74^ and implicated in regulating SC asymmetric cell division^75^ — is expressed from the Xi in SCs (**Figure 4E**). XCI escapees in SCs and myoblasts are located across the length of the X chromosome (**Figure S1I**) and exhibit a higher ratio of Xi:Xa expression compared to genes subject to silencing (**Figure 4F, G**).

**Figure 4.**
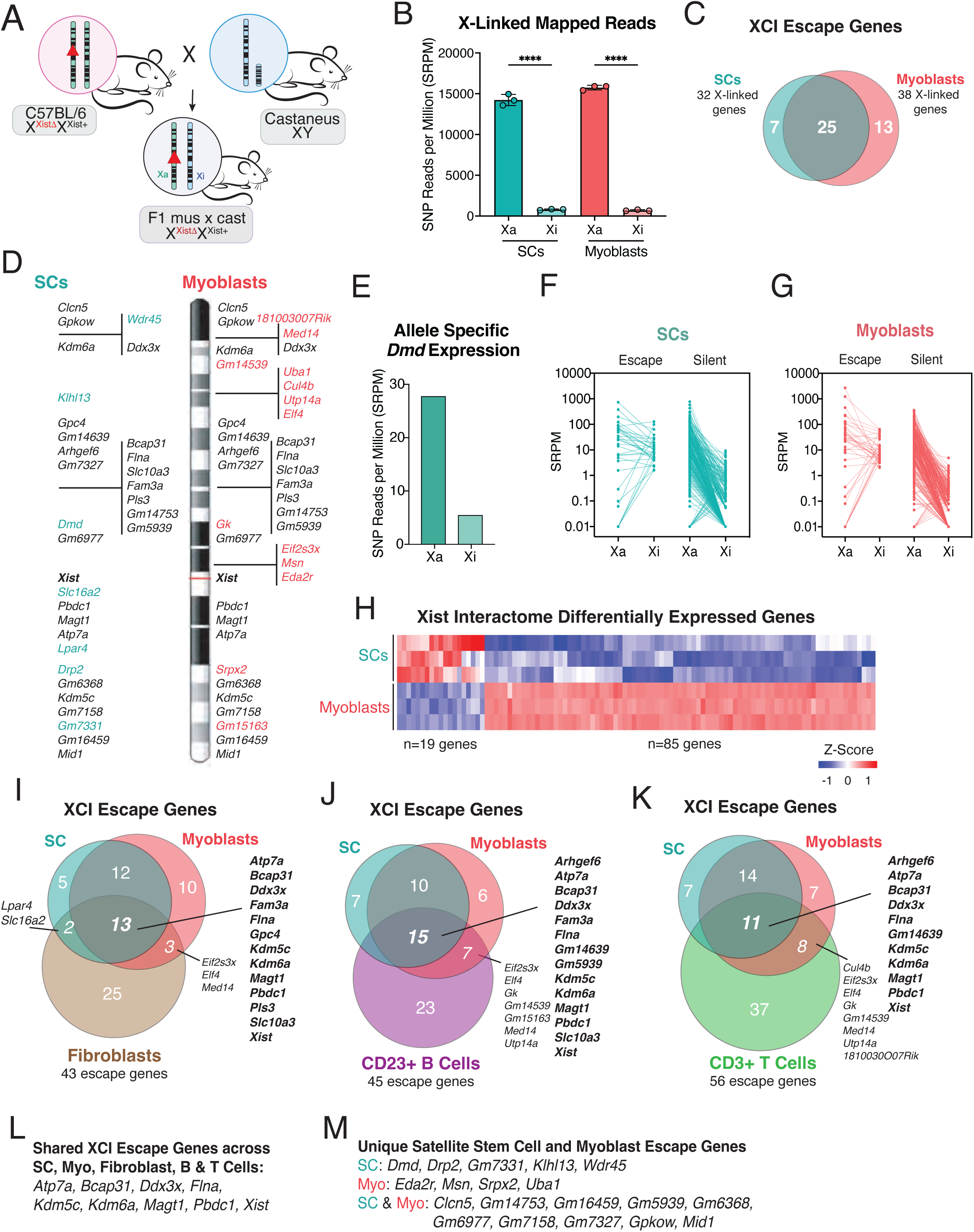
Allele-specific quantification of X-linked genes in SCs and myoblasts reveals muscle-specific genes expressed from the Xi. **A.** Schematic for generating female F1 hybrid mice with skewed XCI for isolation of SCs and myoblasts. **B.** SNP reads per million (SRPM) of X-linked genes expressed from the active (Xa) and inactive (Xi) X chromosomes in SCs and myoblasts (n=3 female mice). **** *p* < 0.0001 by unpaired t test. **C.** Venn diagram of XCI escape genes unique to SCs (n=7 genes) or myoblasts (n= 13 genes) and shared by both cells (n=25 genes). **D.** Location of XCI escape genes along the X chromosome in SCs (left) and myoblasts (right). Shared XCI escape genes across both cell types in black, SC unique escapees in teal, myoblast unique escapees in coral. **E.** Allelic expression of *Dmd*, shown as SRPMs from the Xa or Xi chromosome in SCs. **F-G.** SRPM values for XCI escape and silent genes, showing expression from the Xa and Xi, for SCs (E) and myoblasts (F). **H.** Heatmap of Xist RNA ‘Interactome’ genes that are significantly differentially expressed in SCs and myoblasts. About 100 Xist Interactome genes are expressed in SCs and myoblasts, and cluster into 2 groups. **I.** Comparison of XCI escapees across SCs, myoblasts, and fibroblasts. 13 X-linked genes are shared across 3 cell types and listed in bold**. J.** Comparison of XCI escapees across SCs, myoblasts, and naïve CD23+ B cells. 12 X-linked genes are shared across 3 cell types and listed in bold. **K.** Comparison of XCI escapees across SCs, myoblasts, and unstimulated CD3+ T cells. 11 X-linked genes are shared across 3 cell types and listed in bold. **L.** List of XCI escapees expressed in SCs, myoblasts, fibroblasts, B cells, and T cells. **M.** List of XCI escapees specific to SCs, myoblasts, or both muscle progenitors.

Pathway analysis of XCI escape genes revealed enrichment of pathways including synapse organization, apoptosis, ribonucleoprotein complex biogenesis, and transport of small molecules (**Figure S1J-K**). Several X-linked genes that escape XCI in both SCs and myoblasts have tissue specific functions, as *Fam3a* promotes muscle differentiation^76^, and its expression from the Xi may enhance the transition from SCs to myoblasts. *Flna* inhibits skeletal muscle senescence^77^, and *Magt1* encodes for a magnesium transporter^78^ that regulates magnesium flux which is critical for muscle differentiation^79,80^. These genes highlight the potential physiologic role for XCI escape in muscle, where X-linked genes important for muscle regeneration and homeostasis are expressed from the Xi and Xa in SCs and myoblasts.

Xist RNA binds to various RNA binding proteins for Xist RNA localization and gene silencing^4,81^. To determine whether some of these factors are differentially expressed during SC differentiation into myoblasts, potentially contributing to changes in Xist RNA localization, we examined expression of the ‘Xist RNA Interactome’^13^ genes in SCs and myoblasts. Of the 303 Xist RNA Interactome genes, 285 were expressed in both SCs and myoblasts (**Figure S1L**). We found that 104 Xist RNA Interactome genes were significantly differentially expressed between SCs and myoblasts (**Figure 4H**), and the majority of these (85 genes; 82%) were upregulated in myoblasts. When comparing the genome-wide expression profiles for SCs and myoblasts, we found similar numbers of differential expressed genes for myoblasts and SCs (**Figure S1M**). This finding highlights a potential mechanism where some Xist RNA interacting genes are upregulated during differentiation from SCs to myoblasts, possibly functioning in Xist RNA tethering or recruitment to the Xi.

Finally, we asked how XCI escape genes vary in SCs and myoblasts compared to lymphocytes, (which exhibit dynamic Xist RNA localization) and mouse embryonic fibroblasts (with static Xist RNA localization). We compared lists of XCI escape genes from fibroblasts (n=43 escape genes)^82^ (**Figure 4I**), CD23+ B cells (n=45 escape genes)^23^ (**Figure 4J**), and CD3+ T cells (n=56 escape genes)^13^ (**Figure 4K**). We found that 9 X-linked genes escaped in all cell types measured (**Figure 4L**), including XY gene pairs (*Ddx3x Eif2s3x*, *Kdm5c*, and *Kdm6a*) which are constitutive escape genes in various tissues^16^. We observe that there are fewer overlapping XCI escape genes between muscle cells and the other somatic cells than the cell specific genes, likely reflecting cell type specificity of XCI escape genes that are lost when examining whole tissues. We found that some genes escape XCI exclusively in muscle cells (**Figure 4M**), including *Dmd* and the dystrophin homologue *Drp2*^83^. X-linked *Eda2r* and *Msn* genes are also expressed from the Xi in myoblasts and their upregulation has been implicated in diseases of the skeletal muscle and myopathies^84,85^. Thus, SCs and myoblasts exhibit cell type specific expression of XCI escape genes relevant for muscle physiology, which may contribute to sex differences with muscle regeneration and function.

### SC-specific deletion of Xist does not impact muscle regeneration after injury

Based on our observations of minimal Xist RNA localization and muscle-specific X-linked genes expressed from the Xi in SCs, we asked whether Xist RNA expression would impact skeletal muscle regeneration. We deleted *Xist* in SCs by crossing mice with loxP sites flanking the transcription start site and first 3 exons of *Xist* (“Xist2lox”)^86^ with a tamoxifen inducible Pax7 driven Cre (“Pax7CE”)^87^ and intercrossed positive animals to generate conditional *Xist* homozygous null female animals (Xist SC cKO/cKO; “Xist SC cKO” **Figure S2A, S2B**). Female Xist +/+, Xist SC cKO/+, and Xist SC cKO mice were treated with tamoxifen daily for 5 consecutive days to induce Cre recombinase-mediated loxP recombination in SC nuclei, followed by chemical injury of the right tibialis anterior (TA) muscle via intra-muscular barium chloride injection (**Figure S2C**). Three weeks post injury, TA muscles were isolated and sectioned for hematoxylin and eosin (H&E) staining and immunofluorescence (IF) detection of SCs using anti-PAX7. We quantified the numbers of SCs identified by anti-PAX7 IF within the regenerated regions identified by myofibers with centrally localized nuclei (**Figure S2C** white arrows). We detected no difference in the average number of SCs per area in regenerated TA muscles across all genotypes suggesting that SC self-renewal is not compromised in the absence of *Xist* (**Figure S2E**). Robust myotoxin-induced regeneration was evidenced by the presence of myofibers with centralized nuclei in skeletal muscle cross-sections of all injured mice (**Figure S2E** black arrows). Quantification of muscle morphology after response to injury, either by muscle fiber size or diameter, demonstrated no significant differences between controls and mice lacking *Xist* specifically in SCs (**Figure S2G, H**). We conclude that *Xist* is dispensable in SCs for injury-induced muscle regeneration in mice.

## DISCUSSION

Cytological enrichment of epigenetic features of the Xi exhibit diversity across female immune cells^6,11,12^, where detection of Xist RNA and heterochromatic histone tail modifications with the Xi often correlate with active NF-κB signaling^13^. Here we asked whether active NF-κB signaling correlates with Xist RNA clouds in female adult stem cells, progenitor cells, and terminally differentiated cells from the intestine, blood, and muscle. We find that intestinal progenitor cells exhibit variable Xist RNA localization patterns despite known activated NF-κB signaling status, yet blood progenitors and neutrophils have Xist RNA localization at the Xi that correlates with activated NF-κB. Unlike blood and intestinal cells, stem and progenitor cells in muscle have Xist RNA clouds and pinpoints in the absence of NF-κB activation. While the distribution of Xist RNA patterns changes with age, the Xi in muscle progenitors is dosage compensated yet expresses muscle-specific X-linked genes. Our experiments demonstrate that female somatic cells exhibit variable patterns of Xist RNA localization independent of NF-κB activation status, and that Xist RNA localization changes with cellular stimulation, injury, or differentiation. Thus, our findings indicate that female somatic cells utilize diverse epigenetic mechanisms for XCI maintenance to ensure selective Xi gene expression across a mostly silent chromosome.

We were surprised to find that intestinal progenitor cells at the +4 position of the crypt have significantly reduced frequencies of Xist RNA clouds compared to all other positions within this compartment. Because cells at the +4 position of the crypt are typically quiescent, exhibit slow cell cycle rates, and are considered to be facultative stem cells^50,52–54^, we hypothesized that Xist RNA localization in female ISCs might be influenced by cell cycle rates. Examination of slow-cycling and active cycling stem cells using HopX^88^, Lgr5^56^, and Chga^57,58^ markers revealed no significant differences with frequencies of Xist RNA clouds across reporter positive and negative mice, suggesting that Xist RNA localization is not affected by cell cycling rates for intestinal progenitors. Enteroendocrine cells expressing Chga are known to contain facultative stem cells capable of re-entering the cell cycle following injury^57,58^, and these cells predominantly lacked Xist RNA clouds. Enteroendocrine cells have various toll-like receptors (TLRs) and can secrete cytokines after TLR activation, which also increases NF-κB signaling^89^. It is possible that female enteroendocrine cells mostly lack Xist RNA clouds because of low levels of NF-κB signaling, and that TLR stimulation would increase the frequency of Xist RNA clouds in enteroendocrine cells, as observed for female lymphocytes^11,12^. Additional experiments that quantify features of NF-κB activity and downstream signaling pathways across crypt cell populations would reveal the pathways which influence Xist RNA localization to the Xi in ISCs.

Our results suggest that cell division does not influence Xist RNA localization patterns in immune cells. We did not observe differences with frequencies of Xist RNA clouds and dispersed Xist RNA patterns between slow-cycling HSCs (Lin^-^Sca-1^+^cKit^+^CD150^+^CD48^-^) and fast-cycling MPPs (Lin^-^Sca-1^+^cKit^+^CD150^-^Cd48^-^)^61^, and both HSCs and MPPs predominantly have Xist RNA signals and low frequencies of nuclei lacking Xist RNA. Most B cell progenitors in the bone marrow do not undergo cell division, with the exception of pre-B cells that undergo multiple rounds of cell division, yet all B cell progenitors lack detectable Xist RNA signals^11,90^. Some T cell progenitors also lack Xist RNA localization by RNA FISH,^12^ and these subsets exhibit rapid proliferation. Our study found that post-mitotic neutrophils lack Xist RNA localization, but PMA stimulation for 4-6 hours, which activates NF-κB signaling^64^ without inducing cell division, results in dispersed Xist RNA patterns. We conclude that Xist RNA localization at the Xi for immune cells likely requires active NF-κB signaling, and additional experiments examining Xist RNA localization patterns for other innate immune cell populations are necessary.

Female MEFs have bright Xist RNA clouds on the Xi, and we hypothesized that serum starvation might alter Xist RNA localization patterns in these cells. Serum starvation blocks fibroblast cell division and rapidly reduces levels of nuclear p65, the NF-κB subunit, which reduces NF-κB signaling activity^67^. We were surprised that serum starvation for 2-4 days did not reduce Xist RNA clouds in female MEFs, nor increase the frequency of nuclei lacking Xist RNA signals. It is possible that increased culture time without serum is necessary to affect Xist RNA localization in female MEFs. Another possibility is that NF-κB activation is not required for maintaining Xist RNA localization in MEFs. Myoblasts differentiated from SCs have bright Xist RNA clouds like fibroblasts, consistent with activation of canonical NF-κB signaling in proliferating myoblasts^91^. However, SCs exhibit 4-5 different types of Xist RNA localization patterns which vary in distribution with age. The predominant Xist RNA localization pattern for SCs from ‘young’ mice (∼1 week old) is Xist RNA clouds together with dispersed patterns, yet for adult mouse SC Xist patterns are either pinpoints or absent. Yet SCs from female mice of advanced ages (>2yrs) exhibit predominantly Xist RNA clouds, along with ∼20% of other 4 patterns. Intriguingly, there is dynamic NF-κB activity through both the canonical and alternative signaling pathways, in skeletal muscle from birth into adulthood and with advanced age. There is high NF-κB activity in skeletal muscle during the first 2 weeks after birth, with elevated protein levels of p65 and p50 subunits of NF-κB, and levels decline at 3 weeks into adulthood, yet this activity was localized to stromal fibroblasts and not SC progenitors^92^. It is unclear whether NF-kB activation in stromal fibroblasts influences NF-κB activity and perhaps Xist RNA localization in SCs in post-natal day 7 mice, and whether NF-κB activation levels contribute to the abundance of SC nuclei lacking detectable Xist RNA signals (40-100%) in adult mice. Previous work found increased NF-κB activity in aged (∼2yr) skeletal muscle, where increased NF-κB signaling in myofibers and myotubules reduces SC function^93^. It is likely that age-induced inflammation and elevated NF-κB activity in skeletal muscle contributes to the high frequency of Xist RNA clouds in aged SCs. Additional work is necessary to determine how stromal fibroblasts and the stem cell niche influence NF-κB levels in SCs over the lifespan, and whether NF-κB levels modulate each type of Xist RNA localization pattern in SCs.

There is a diversity of Xist RNA deletion phenotypes in mice, which vary from embryonic lethality to mild/no effects, depending on the tissue and timing of deletion^94^. Given the variation of Xist RNA localization patterns in SCs, we deleted *Xist* in SCs to determine whether SC-dependent regeneration would be affected. Muscle tissue injury results in rapid activation and differentiation of SCs into myoblasts, which then further differentiate to replenish muscle fibers^95^. *Xist* deletion in SCs did not affect survival or lifespan (data not shown) and did not impact numbers of SCs within muscle fibers nor SC regeneration efficiency after barium chloride injection. One possibility is that *Xist* deletion in combination with advanced age or in mice lacking telomerase with shorter telomeres^96^ may impair SC mediated repair, as muscle repair potential is compromised with age and associated with muscle inflammation and reduced SC function^97^. Alternatively, it is possible that muscle injury induces upregulated expression of X-linked genes important for SC function and myogenesis such as *Dmd*, *Flna*, *Fam3a*, and *Magt1* to be expressed from both the Xa and Xi, and because these genes ‘escape’ XCI at steady state in SCs and myoblasts, *Xist* deletion and injury together could increase their expression from the Xi. Deletion of *Xist* can reduce enrichment of H3K27me3 and H2AK119-Ub modifications across the Xi in somatic cells^98^, which would facilitate less chromatin compaction and accessibility for transcriptional activators. Additional work is necessary to determine the fidelity of Xi gene repression following *Xist* deletion in SCs, and whether there are changes with DNA methylation levels at CpG regions of Xi promoters or with enrichment of histone variant macroH2A and other heterochromatic histone modifications across the Xi.

Quantification of allele-specific expression of X-linked genes in SCs and myoblasts revealed that ∼11-13% of expressed genes are expressed from the Xi as well as the Xa. These XCI escapees are mostly similar between SCs and myoblasts and include several genes important for muscle function and regeneration. Strikingly, among the few genes we identified as being transcribed from the Xi in SCs is *Dmd*. It is possible that *DMD* will also escape XCI in human SCs, as the human Xi exhibits more XCI escape genes compared to the mouse chromosome^99^. Using GTEx datasets surveying diverse tissues, *DMD* exhibits significant female-biased expression in three tissues, yet did not exhibit detectable biallelic expression^18^, likely because specific cell types were not examined. While most female carriers of Duchenne Muscular Dystrophy (DMD) remain clinically unaffected, approximately 3–8% present as manifesting carriers, displaying skeletal muscle dysfunction that ranges from mild proximal weakness to severe gait impairment. Skewed XCI has long been proposed as a determinant of variable dystrophin expression in carriers^100^, however, our findings raise the possibility that heterogeneity in XCI escape at the *DMD* locus in human SCs may represent an additional mechanism contributing to the diverse clinical phenotypes observed in this population.

While we predicted that X-linked genes with Y-chromosome homolog would be shared across all cell types (*Ddx3x, Kdm5c, Kdm6a, Flna, Pbdc1),* we were surprised to find several shared escape genes lacking a Y-homolog (*Atp7a, Bcap31, Magt1*). XCI escape genes usually exhibit sex-biased gene expression^101^, and skeletal muscle regeneration exhibits sex differences. Females have fewer SCs yet exhibit faster recovery following barium chloride induced muscle injury, where females have increased muscle force and fiber size post-recovery compared to males^102^. Sex differences with SC function and muscle regeneration are likely influenced by estrogen and other sex hormones^103^, yet our study suggests that XCI escape genes likely contribute to the observed sex differences with SCs and responses to injury. One prediction is that muscle injury will induce increased expression of muscle-related X-linked genes from the Xi in SCs and myoblasts, and perhaps reactivate additional genes from Xi that are silent at steady state. Additional work is needed to determine which X-linked genes exhibit sex-biased expression patterns in SCs and myoblasts, and how these genes contribute to observed sex differences with muscle repair following injury. Taken together, our findings highlight the diverse and complex mechanisms that govern XCI maintenance and X-linked gene expression across female somatic cells. Contributions from the X chromosome likely contribute to sex bias in a wide variety of tissue function in physiology and disease, and dissecting epigenetic mechanisms that control X-linked gene dosage may inform innovations in sex-specific treatments.

---

**Supplemental Figure 1.**
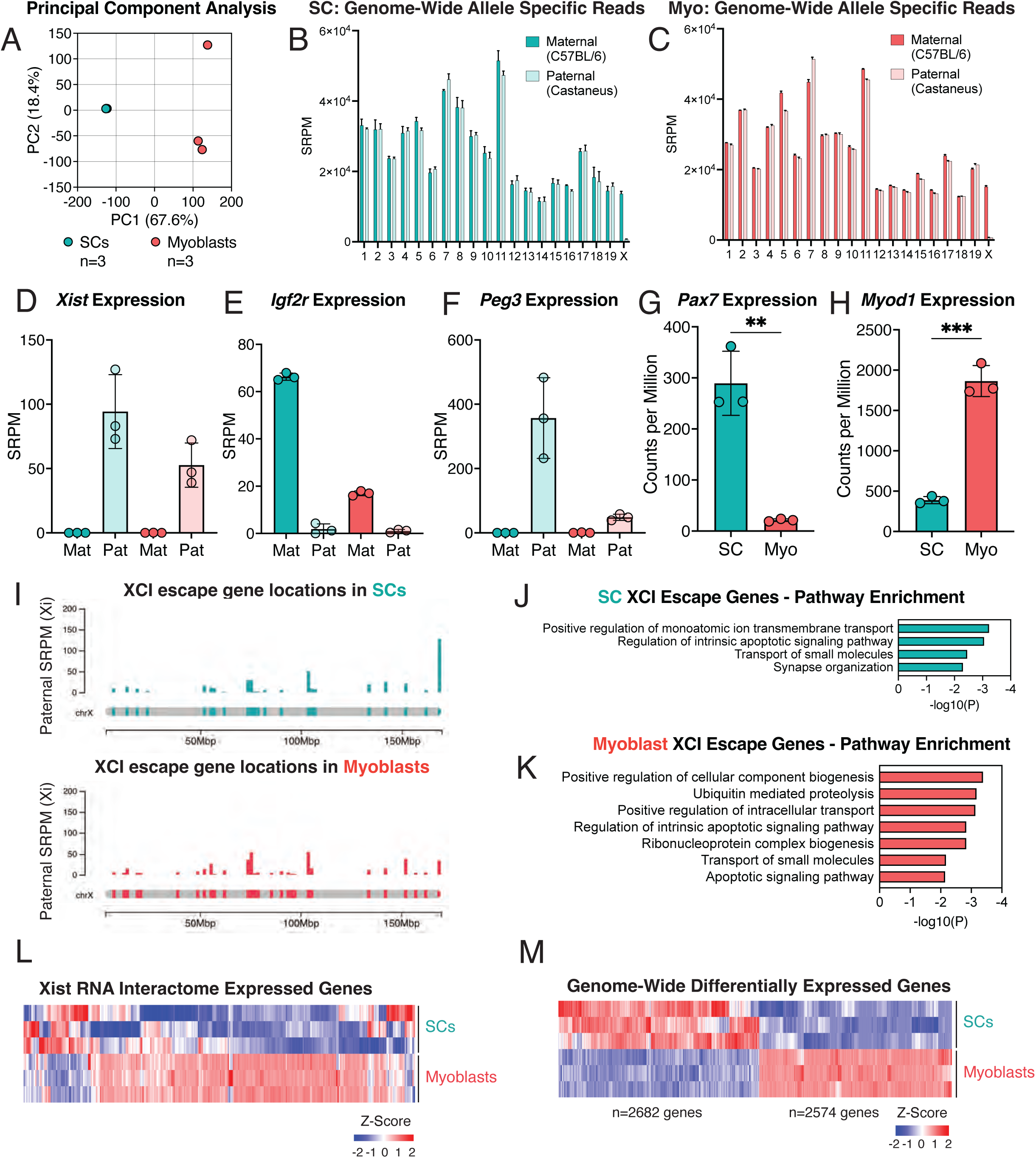
Allele specific profiling of SCs and myoblasts from female F1 mus x cast mice. **A.** Principal component analysis of RNAseq from n=3 SCs and n=3 myoblasts from female F1 mice. **B-C.** SNP reads per million (SRPM) originating from the maternal (C57BL/6) or paternal (Castaneus) chromosomes across all autosomes and the X chromosome in SCs (**B**) and myoblasts (**C**). Only the X chromosomes show significant expression differences between alleles. **D-F.** Verification of allelic expression of X-linked (*Xist;* **D**) and autosomal (*Igf2r;* **E***, Peg3;* **F**) imprinted genes. SRPMs from the maternal (C57BL/6) or paternal (Castaneus) chromosome for each gene in SCs and myoblasts. **G.** Expression of *Pax7* in SCs and myoblasts, shown as counts per million. ** *p* < 0.01 by unpaired t test. **H.** Expression of *Myod1* in SCs and myoblasts, shown as counts per million. *** *p* < 0.001 by unpaired t test. **I.** Location of XCI escape genes along the X chromosome in SCs (top) and myoblasts (bottom). Height of bars represents SRPM for X-linked genes. **J.** Pathways for XCI escape genes in SCs using Metascape. **K.** Pathways for XCI escape genes in myoblasts using Metascape. **L.** Heatmap showing expression levels for all Xist RNA Interactome genes (n=285) in SCs and myoblasts. About ∼100 Xist Interactome genes are expressed in SCs and myoblasts (see Figure 4G). **M.** All differentially expressed genes (n=5256), genome-wide, comparing SCs and myoblasts.

**Supplemental Figure 2.**
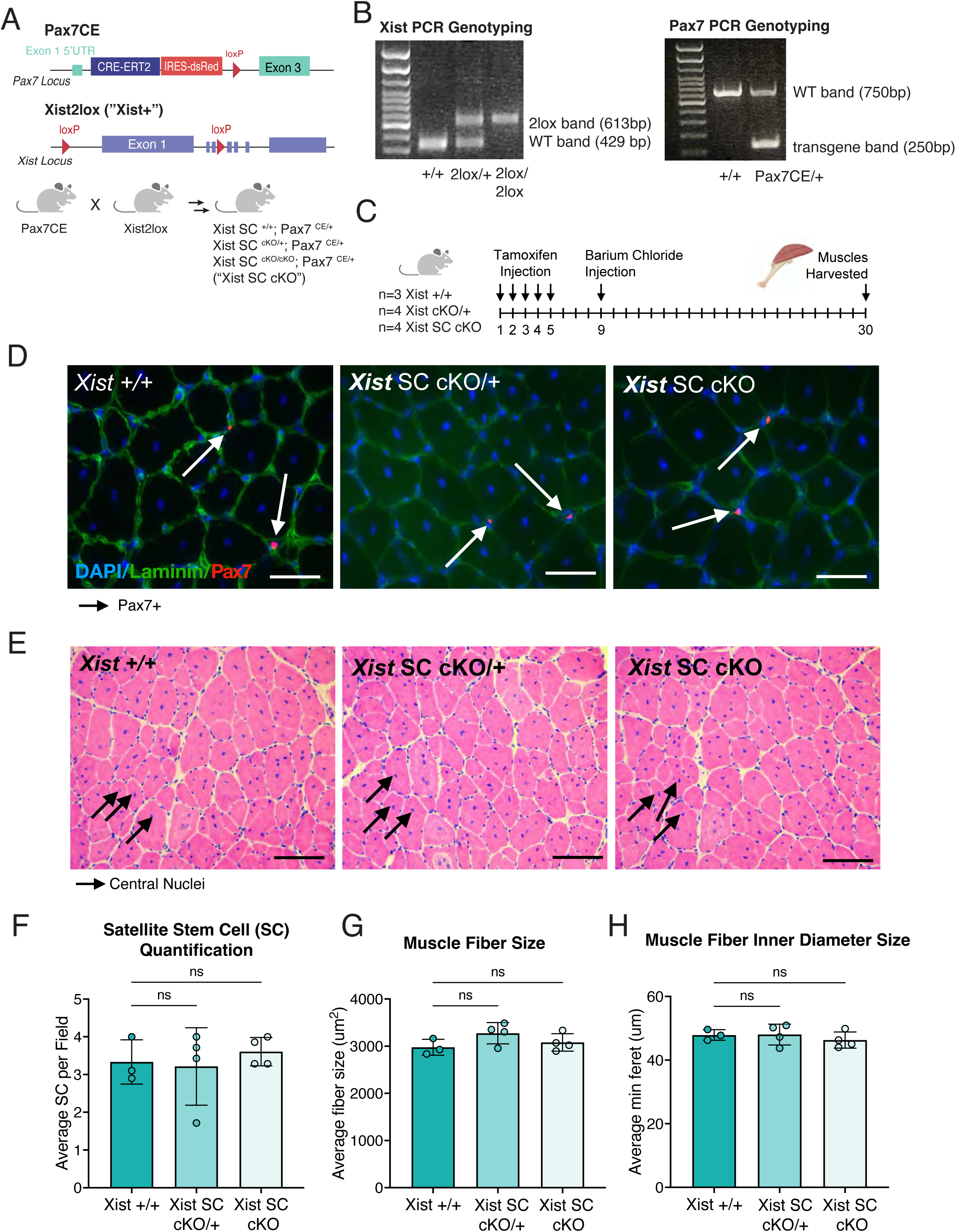
X*i*st deletion does not alter SC-mediated muscle regeneration after chemical injury. **A.** Breeding scheme to generate mice with a heterozygous tamoxifen-inducible heterozygous (“Xist +/flox”) or homozygous (“Xist flox/flox”) loss of *Xist* in Pax7+ SCs. **B.** Representative genotyping PCRs for *Xist* deletion (left) and detection of Pax7 CRE-ERT2 transgene (right) of female Xist cKO/+, Xist cKO/cKO, and wildtype mice for muscle injury experiments. Amplicon sizes for wildtype (WT) and transgenes are shown next to corresponding band. **C**. Experimental design utilized to induce *Xist* deletion by tamoxifen injections for 5 days, followed by barium chloride injection at day 9 to induce muscle injury, and muscle harvest at day 30. **D.** Representative images of muscle fibers stained by immunofluorescence for laminin (green) and Pax7 (red). White arrows denote Pax7+ SCs. **E.** Representative images of muscle fibers stained with hematoxylin and eosin (H&E). Black arrows denote muscle fibers with central nuclei, a signature of response to injury. **F.** Quantification of the average number of satellite stem cells per field by immunofluorescence in n=3 Xist +/+, n=4 Xist +/-, and n=4 Xist -/- mice after tamoxifen treatment and muscle injury. “Ns” not significant by ordinary one-way ANOVA followed by Dunnett’s multiple comparisons test. **G.** Quantification of the average fiber size (um^2^) of muscles from n=3 Xist +/+, n=4 Xist +/-, and n=4 Xist -/- mice after tamoxifen treatment and muscle injury. “Ns” not significant by ordinary one-way ANOVA followed by Dunnett’s multiple comparisons test. **H.** Quantification of the average inner diameter (um) of muscle fibers from n=3 Xist +/+, n=4 Xist +/-, and n=4 Xist -/- mice after tamoxifen treatment and muscle injury. “Ns” not significant by ordinary one-way ANOVA followed by Dunnett’s multiple comparisons test.

## EXPERIMENTAL PROCEDURES

### Resource availability

Lead contact: For additional information, please contact Montserrat Anguera.

Materials availability: New reagents were not developed in this study.

Data and code availability: All sequencing data generated in this study has been deposited to the NCBI GEO database. Access data using the accession number GSE306504.

### Mice

Female C57BL/6 mice (aged 2-6 months) were purchased from the Jackson Laboratory and used for whole intestines, HSCs, neutrophils, and satellite cell isolations. For sorted intestinal populations, mice were obtained from the Jackson Laboratory: Hopx-CreERT2 (#017606), *Lgr5^EGFP-IRES-CreERT2^*(#008875). For Hopx Tomato+/-cell isolation, we used female *Hopx^ERCre/+^; R26^LSL-tdTomato^* ^55^ mice. The Chga reporter mice (Chga^CreERT2-2A-tdTomato^) were generated by C. Lengner^57^, and we were limited by availability of female mice containing the Chga marker as this mouse line was lost during this study. Pax7*-*Avi−2A-GFP mice – for equimolar expression of both PAX7 and membrane-bound GFP (from the *Pax7* locus via 2A peptide cleavage) – were used for isolation of Pax7+ (GFP+) SCs and will be described by Dr. Lepper in a separate study. Pax7Cre-ERT2; Xist2lox mice were generated by multi-generation crosses of Pax7Cre-ERT2 (“Pax7CE”)^104^ and Xist^2lox/2lox^ mice^105^ to generate WT controls (Pax7CE+;Xist+/+), heterozygous (Pax7CE+;Xist^+/fl^), and homozygous (Pax7CE+;Xist^fl/fl^) animals. *Xist* deletion and presence of Pax7 Cre-ERT2 was confirmed using PCR genotyping of genomic DNA from mouse tails (**Figure S1B)**. F1 mus x cast mice with skewed XCI were generated by mating Xist^+/fl^ C57BL/6 females with wild-type *Mus Castaneus* (Cast) males. F1 Xist^+/fl^ females from this mating always inactivate the paternal castaneus X chromosome. All mice were maintained at the Penn Vet animal facility, and all experiments were approved by the University of Pennsylvania Institutional Animal Care and Use Committee (IACUC).

### Generation of intestinal tissue sections for Xist RNA FISH

Duodenum from female mice were harvested, cut into segments and flushed with 1x PBS followed by 3.7% formaldehyde on ice. Segments were fixed in 3.7% formaldehyde for 3 hours, then equilibrated overnight in sucrose-formaldehyde solution. Segments were then flash-frozen in OCT using 2-methyl butane and sectioned into 5-micron slices (1-1.5 cell width).

### Intestinal crypt cell isolation

The gastrointestinal tracts of Hopx-CreER, Lgr5-GFP, and Chga-CreERT2-2A-tdTomato mice were dissected and the most proximal 15cm of the small intestine was isolated in PBS. The tissue was washed in fresh PBS, splayed open, and villi were scraped off using a coverslip. The tissue was transferred to a tube containing 10ml of 1X HBSS with 1mM NAC and 10mM EDTA and was placed on a rotator at 4°C for 45 minutes. After the incubation, the tissue was vortexed for 30 seconds followed by a 30 second rest period on ice; this was performed three times. After vortexing, the tissue was filtered through a 70uM filter and centrifuged at 300g for 3 minutes. To generate a single cell suspension, the cell pellet was resuspended in buffer containing DNAse (35ug/mL) and Liberase (20ug/mL) and incubated at 37°C for 20 minutes. Following digestion, the cells were washed in PBS and resuspended in FACS buffer (1% BSA in PBS) prior to sorting. Cells were sorted at the CHOP Flow Cytometry Core using the FACSJazz Sorter or the MoFlo Astrios Sorter. Cells were sorted on either tdTomato expression (Hopx+ or Chga+) or GFP expression (Lgr5+). Sorted cells were immediately cytospun on slides, fixed in 4% paraformaldehyde, and processed for Xist RNA FISH.

### Hematopoietic stem cell isolation

Bone marrow from female mice was harvested by flushing femurs and tibiae with 1% FBS in PBS on ice. Bone marrow was first lineage depleted using the Direct Lineage Cell Depletion Kit (Miltenyi Biotec 130110170). Cells were stained with the following antibodies from BioLegend: Ter119 (clone TER-119), B220 (clone RA3-6B2), CD19 (clone 6D5), CD3e (clone 17A2), TCRb (clone H57-597), CD8a (clone 53-6.7), NK1.1 (clone PK136), CD11b (clone 6D5), CD11c (clone N418), Gr1 (clone RB6-8C5), Sca-1 (clone D7), cKit (clone 2B8), CD150 (clone TC15-12F12.2), and CD48 (clone HM48-1). Long-term hematopoietic stem cells (LT-HSC) and multipotent progenitors (MPP) were sorted at the Penn Cytomics and Cell Sorting Resource Laboratory on a BD FACS Aria II using the following strategy: Lineage (Lin): Ter119+ B220+ CD19+ CD3e+ TCRb+ CD8a+ NK1.1+ CD11b+ CD11c+ Gr1+; LT-HSC: Lin-Sca-1+ cKit+ CD150+ CD48-; MPP: Lin-Sca-1+ cKit+ CD150-Cd48-. Sorted cells were immediately cytospun on slides, fixed in 4% paraformaldehyde, and processed for Xist RNA FISH.

### Neutrophil isolation

Bone marrow from female mice was harvested by flushing femurs and tibiae with ice-cold 1% FBS in PBS. Neutrophils were isolated from bone marrow using a neutrophil enrichment kit (Miltenyi Biotec 130097658). Cells were cultured in RPMI-1640 (Invitrogen 11875085) containing 10% FBS and stimulated with 100nM PMA. Cells were harvested for slide preparation at 0-, 2-, 4-, and 6-hours post-stimulation. Cells were cytospun on slides, fixed in 4% paraformaldehyde, and processed for Xist RNA FISH.

### Induction of quiescence for MEFs using serum starvation

Primary mouse embryonic fibroblasts (MEFs) were isolated as previously described^106^. Ears from female mice were cut into 3mm size pieces then incubated in collagenase D-pronase solution for 90 minutes at 37°C with agitation, then ground and filtered through a 70um cell strainer into complete medium (RPMI with 10% fetal calf serum, 50uM 2-mercaptoethanol, 100uM asparagine, 2mM glutamine, 1% penicillin-streptomycin). Cells were washed then plated with complete medium supplemented with 10ul amphotericin B (250ug/ml stock), and cultured until they reached 70% confluency. Fibroblasts were split and plated in normal media (10% FBS (Hyclone SH30071.03), 1% Pen/Strep (Invitrogen 15140122), 1% Glutamax (Invitrogen 35050061) in DMEM (Invitrogen 11965084)) and allowed to adhere to the plate, then switched to low serum media (0.02% FBS, 1% Pen/Strep, 1% Glutamax in DMEM). After 2 or 4 days of culture, cells were collected, cytospun onto slides, fixed in 4% paraformaldehyde, and processed for Xist RNA FISH.

### Muscle SC and myoblast isolation

SCs were isolated as previously published^107^. Hind limb muscles were dissected from female mice, then minced, then incubated in muscle dissociation buffer (700-800U/ml collagenase II in Hams F-10 supplemented with 10% horse serum and 1x penicillin-streptomycin) in a 37°C shaking water bath for 1 hour. Cells were spun down and resuspended in collagenase II and 11U/ml dispase solution and pipetted roughly 15 times. Cells were incubated in 37°C shaking water bath for 30 minutes, then the digested muscle samples were passed through a 30ml syringe with a 20-gauge needle ∼10 times. Cells were spun down, resuspended, and strained through a 40um nylon cell strainer and Pax7*-*Avi−2A-GFP samples were gated for GFP+ fraction. SC samples for chromosome discrimination RNA-Seq were isolated via negative and positive cell surface marker expression using antibodies from BioLegend: CD31 (clone MEC13.3), CD45 (clone 30-F11), Sca1 (clone D7), VCAM1 (clone 429). SCs were sorted as CD31-CD45-Sca1-, VCAM1+. A proportion of sorted SCs were cultured for two days to stimulate activation and differentiation into myoblasts in 2-well slide chambers prior to RNA FISH.

### Xist RNA FISH on cryo-fixed intestinal sections

Duodenum sections were fixed in 4% PFA for 20 mins, rinsed twice in 1X PBS under gentle agitation, and permeabilized 1h in 70% ethanol at 4°C. Sections underwent two 5 minute washes in wash buffer (10% formamide, 1X SSC) then incubated overnight with Xist RNA probes as previously described^10,11^. Sections were then incubated in wash buffer for 30 minutes in a humidified chamber, then DAPI-stained and mounted. Images were obtained using a Nikon Eclipse microscope.

### Sequential Xist RNA fluorescent in situ hybridization (FISH) and immunofluorescence (IF)

Xist RNA FISH was performed as previously described^10,11^. Briefly, 2 Cy3-labeled 20-nucleotide oligo probes were designed to recognize regions within exon 1 of Xist (synthesized by IDT). Sequential immunofluorescence for histone marks was performed using a primary antibody for H2AUb119K (Cell Signaling 8240S) and a secondary AF488-conjugated goat anti-rabbit IgG antibody (Abcam ab150077-500UG). Samples were imaged using a Nikon Eclipse microscope.

### Muscle injury experiments

Female Pax7CE; Xist SC cKO mice and controls (3 months of age) were IP injected with 3mg of tamoxifen for 5 consecutive days then left to recover for 3-5 days. Mice were sedated, shaved, and injected with 50ul of 1.2% barium chloride (Sigma-Aldrich 202738) into the right TA muscle to stimulate injury^108^. After recovery (21 days post-injection), animals were sacrificed, the TA muscle was removed, mounted on cork, covered in OCT, and cryo-frozen using super-chilled 2-methyl butane^109^.

### H&E staining of muscle sections

Histology of muscle sections was performed as previously described^109^. Briefly, muscles were sectioned at 10um thickness and collected on Histobond slides (Azer Scientific UNI75251+). Samples were fixed in 100% ethanol, incubated for 4.5min in Gill’s II hematoxylin, then treated with Scott’s tap water for 10sec. Samples were stained with Eosin for 45sec before destaining using ethanol and Histoclear 1/2. Tissue was mounted using Cytoseal-60 and imaged using bright-field microscopy using a ToupCam camera (UCMOS05100KPA).

### IF and scoring of muscle tissue sections

Flash frozen muscle was cryo-sectioned at 10um thickness and collected on Histobond slides (Azer Scientific UNI75251+). IF was performed as previously described^109^. Briefly, tissue was fixed in 4% paraformaldehyde/PBS and permeabilized using 0.3% Triton-X/PBS. Samples were blocked first with M.O.M. blocking reagent (Vector Laboratories BMK-2022), then with goat blocking solution (1% Blocking Powder (Perkin Elmer, FP1012)/10% goat serum in 0.05% Triton-X/PBS). Sections were incubated in primary antibodies overnight at 4°C, followed by incubation in secondary antibodies for 1 hour at room temperature. Antibodies used were as follows: anti-Laminin (Sigma L9393; 1:1000), anti-Pax7 (Developmental Studies Hybridoma Bank; 1:5), goat anti-rabbit IgG Alexa488 (Invitrogen A11034; 1:1000), and anti-mouse IgG1 Alexa568 (Invitrogen A-21124; 1:1000). All quantification was performed blinded. Using ImageJ software, average fiber size and minimum Feret were quantified through analysis of anti-Laminin and DAPI overlays in SMASH^110^. Quantification of SC number was performed using overlays of anti-Laminin, anti-Pax7, and DAPI in ImageJ software. Images were taken at 20x magnification, and sub-laminal Pax7+ SCs were manually counted.

### Allele-specific RNA sequencing

SCs from skeletal muscle of female F1 mus x cast mice were isolated and sorted, then sorted SCs were cultured for two days to stimulate differentiation in myoblasts. SCs and myoblasts were collected in TRIzol (Invitrogen 15596026) and RNA isolation was performed. Libraries were prepared using SMART-Seq HT PLUS (Takara Bio USA, Inc. R400749) with 15 cycles of amplification. Samples were run on the Illumina NextSeq 500 using 75bp single-end reads, yielding between 30 million to 80 million total reads. We quantified allele-specific gene expression for SCs and myoblasts as previously described^13^. We created an N-masked mm10 (C57BL/6j; Ensembl GRCm38) genome using SNPsplit^111^. SNPs were derived from the *Castaneus* genome for the C57BL/6 x Cast F1 mice. The resulting N-masked genomes were used to generate respective STAR (v2.7.1a) indexes. Reads were aligned using STAR with alignEndsType set to EndToEnd, and outSAMattributes set to NH HI NM MD to allow for compatibility with SNPsplit. STAR-aligned files were then passed through SNPsplit for allele-specific sorting of reads. Reads were quantified using featureCounts^112^ on both the diploid N-masked aligned reads and the haploid allele-specific aligned reads. XCI escape genes were defined using 3 thresholds of expression as previously described^16,113^. Briefly, diploid gene expression was first calculated in RPKM (reads per kilobase per million mapped reads), and genes were called as expressed if their diploid RPKM was > 1. For every X-linked gene that passed this threshold, haploid gene expression was calculated in SRPM (allele-specific SNP-containing exonic reads per 10 million uniquely mapped reads), and genes which had a Xi-SRPM > 2 were considered to be expressed from the Xi. Finally, a binomial model estimating the statistical confidence of XCI escape probability was applied to the genes passing the first 2 thresholds. This model compares the proportion of Xi-specific reads to the total Xi + Xa reads and calculates a 95% confidence interval. If the 95% lower confidence limit of a gene’s escape probability was greater than 0, it was called as an XCI escape gene. Our code for calling XCI escape genes (escape_gene_calculator) is publicly available on github (https://github.com/Montserrat-Anguera/escapegenecalculator). XCI escape genes must meet all three thresholds for all n=3 biological replicates. The R package chromoMap was used to generate chromosome maps^114^. To graph the log SRPM of Xa vs Xi transcripts an arbitrary value of 0.01 was added to all SRPM values. For differential gene analysis between SCs and myoblasts, we used the N-masked index created above and aligned using STAR with alignEndsType set to SortedyCoordinate, and quantmode set to GeneCounts. STARaligned files were then quantified using featureCounts. Reads were filtered to have a counts per million (CPM) >1 for at least 3 samples prior to quantile normalization. For PCA analysis, distance was calculated using the dist function (Method = “Euclidean”) and clusters were calculated using the hclust function (Method = “complete”). Differential gene lists were generated using the DeSeq2 package^115^. For Xist RNA Interactome analysis, a list of known binding proteins was compiled manually from published literature^116–119^. This list was filtered to exclude genes that were not found to be differentially expressed between SC and myoblasts. A heatmap of the remaining 104/303 genes was generated using hclust function (Method = “spearman”) and plotted using the heatmap2 function. Metascape was used to perform pathway enrichment on escape genes in SCs and myoblasts^120^.

## ACKNOWLEDGEMENTS

We would like to thank M. Cancro and T. Laufer for critical discussions to conceptualize this study, and L. King for comments and edits to the manuscript. We also thank A. Dubin and A. Driscoll for assistance with genotyping *Xist* and *Pax7* mouse strains. We are grateful to the UPenn NIDDK P30 Digestive and Liver Disease Center for secondary antibodies. This research was supported by NIH R21 AI124084, R01 AI134834, R01 AI168047, and Lupus Research Alliance Target in Lupus award (to M.C.A.), Penn MSTP T32 Training Grant GM07170 (to C.D.L.), the National Institutes of Health NIAID F30 AI174437 (to C.D.L.), and NIH R01 AR078231 (to C.L.).

## AUTHOR CONTRIBUTIONS

M.C.A., C.L., C.D.L., and C.J.L. conceptualized the study. C.D.L., D.S.H., I.S., N.L., A.V. and C.L. performed experiments. C.D.L., D.S.H., and C.L. analyzed data. I.S., C.D.L., and M.C.A wrote the first drafts of the manuscript, and M.C.A. extensively edited all versions of the manuscript. C.D.L, E.M.W., C.L, C.J.L., and M.C.A. edited the final versions of the manuscript. E.M.W. and M.C.A. uploaded the sequencing data and manage data and code availability.

